# A partially sex-reversed giant kelp sheds light into the mechanisms of sexual differentiation in a UV sexual system

**DOI:** 10.1101/2021.02.28.433149

**Authors:** Dieter G. Müller, Enora Gaschet, Olivier Godfroy, Josselin Gueno, Guillaume Cossard, Maritta Kunert, Akira F. Peters, Renato Westermeier, Wilhelm Boland, J. Mark Cock, Agnieszka P. Lipinska, Susana M. Coelho

**Affiliations:** Fachbereich Biologie der Universität Konstanz,78457 Konstanz, Germany; Sorbonne Université, UPMC Univ Paris 06, CNRS, Integrative Biology of Marine Models, Station Biologique de Roscoff, CS 90074, F-29688, Roscoff, France; Department of Bioorganic Chemistry, Max Planck Institute for Chemical Ecology, Jena, Germany; Bezhin Rosko, 29250 Santec, France; Instituto de Acuicultura, Universidad Austral de Chile, Casilla 1327, Puerto Montt, Chile; Max Plank Institute for Developmental Biology, Tübingen, Germany

## Abstract

In UV sexual systems, sex is determined during the haploid phase of the life cycle and males have a V chromosome whereas females have a U chromosome. Previous work in the brown algal model *Ectocarpus* revealed that the V chromosome has a dominant role in male sex determination and suggested that the female developmental program may occur by ‘default’, triggered in the absence of the male master sex determination gene(s). Here, we describe the identification of a genetically male giant kelp strain presenting phenotypic features typical of a female, despite lacking the U-specific region. The conversion to the female developmental program is however incomplete, because gametes of this feminised male are unable to produce the sperm-attracting pheromone lamoxirene. We identify the transcriptomic patterns underlying the male and female specific developmental programs, and reveal the faster evolutionary rates of male-biased genes compared to female-biased and unbiased genes. Moreover, we show that the phenotypic feminisation of the variant strain is associated with both feminisation and de-masculinisation of gene expression patterns. Importantly, the feminisation phenotype was associated with the dramatic downregulation of two V-specific genes including a candidate sex-determining gene on the V-specific region. Our results reveal the transcriptional changes associated with sexual differentiation in a UV system with marked sexual dimorphism, and contribute to disentangling the role of sex-linked genes and autosomal gene expression in the initiation of the male and female developmental programs. Overall, the data presented here imply that the U-specific region in the giant kelp is not required to initiate the female developmental program, but is critical to produce fully functional eggs, arguing against the idea that female is the ‘default’ sex in this species.

## Introduction

Females and males often differ dramatically in appearance and behaviour. These differences, which are referred to as sexual dimorphism, are principally the result of natural and/or sexual selection for traits that influence the fitness of each sex. Genetically, however, females and males are nearly identical differing by only a few genes located on sex-specific chromosomes (such as Y chromosomes in mammals, W chromosomes in birds, or U and V chromosomes in mosses and many algae; Coelho et al. 2018; Umen and Coelho 2019). Consequently, sexually dimorphic traits are to a large extent a result of differential expression of (autosomal) genes that are present in both sexes (Grath and Parsch 2016).

Autosomal sex biased gene expression patterns in diploid XY and ZW systems have been intensively investigated in recent years (reviewed in Grath and Parsch 2016). Such studies have revealed that sex-biased gene expression is abundant in many animal and plant species, although its extent may vary greatly among tissues or developmental stages. In species with genetic sex determination, sex chromosome-specific processes, such as dosage compensation, also may influence sex-biased gene expression. Sex-biased genes, especially genes with male-biased expression in XY systems, often show elevated rates of both protein sequence and gene expression divergence between species, which could have a number of causes, including sexual selection, sexual antagonism, and relaxed selective constraint (Grath and Parsch 2016; Whittle and Extavour 2019). The molecular mechanisms that lead to sex-biased gene expression, however, are yet to be elucidated (Grath and Parsch 2016).

In contrast to the well-studied XY and ZW systems, knowledge about sex-biased gene expression, sexual dimorphism and the link between sex-chromosomes and control of sex-biased gene expression in UV haploid sexual systems remains relatively scarce. UV systems are abundant among Eukaryotes, but are largely understudied (Coelho et al. 2018; Umen and Coelho 2019). In UV systems, sex is determined after meiosis by a male or a female sex-determining region (SDR). After meiosis, if a daughter cell inherits the U-chromosome (containing a U-specific region), it will develop into a female individual (female gametophyte) that at maturity will produce female structures (oogonia) and female gametes (eggs). If the daughter cell inherits a V-chromosome, it will develop into a male individual (male gametophyte), producing male reproductive structures (antheridia), where male gamete cells are produced by mitosis at maturity. Sexual dimorphism can be minor, as in the brown alga *Ectocarpus*, where male and females differ very little, but sexual dimorphism may be more marked in algae such as the giant kelp *Macrocystis pyrifera* (Luthringer et al. 2015; Mignerot and Coelho 2016). The presence of shared orthologues in the SDRs of the Ectocarpales and the giant kelp indicates that the U and V chromosomes is these organisms are derived from the same ancestral autosome (Lipinska et al. 2017).

The SDRs of UV systems contain sex-determining gene(s) that initiate the sexual determination, by regulating sex-specific patterns of expression of downstream effector autosomal genes. In *Ectocarpus*, genetic analysis has shown that the male V-specific region (VSR) is dominant over the female U-specific region, and that the female developmental program is triggered in the absence of the VSR (and presence of the U-specific region) (Müller 1975; Ahmed et al. 2014). These results have led to the idea that a master male sex-determining gene(s) is located on the V-specific region (Ahmed et al. 2014; Lipinska et al. 2017) and that female sex may be initiated ‘by default’, in the absence of the VSR (Ahmed et al. 2014). The U-specific genomic region may therefore not be strictly necessary for initiation of the female developmental program, with female development exclusively relying on autosomal sex-biased gene expression. This idea is further supported by the fact that female-specific genes on the U-specific region present signs of degeneration and relatively low levels of expression (Ahmed et al. 2014; Avia et al. 2018). Hence, an individual with a V chromosome with an impaired male master sex-determining gene(s) would be anticipated to develop into a phenotypic female. A variant or mutant line where the genotypic sex is uncoupled from phenotypic sex would be the ideal system to test this hypothesis and to understand the role of sex-specific and (autosomal) sex-biased genes in the events leading to male and female developmental programs.

We describe here the identification of a variant strain of an organism with UV sex chromosomes, the giant kelp *Macrocystis pyrifera*, that exhibits a range of phenotypic features typical of a female, despite being genetically male. The availability of this feminised line, together with closely related male and female lines, provided access to the molecular events underlying the initiation of the male versus female developmental program in this ecologically important organism and to disentangle the role of sex-linked genes and autosomal gene expression in the initiation of sex-specific development.

We show that a considerable proportion of the transcriptome of the giant kelp is sex-biased, and that the morphological feminisation of the variant line is associated with extensive feminisation and de-masculinisation of autosomal gene expression patterns. Male-biased genes displayed elevated rates of evolution compared to female-biased and unbiased genes, consistent with the notion that sex-specific selection may be stronger in males than females in this species. We show that two genes within the VSR exhibit a significantly reduction in transcript abundance in the feminised line, suggesting their involvement in sexual differentiation. Our observations indicate that the female program in this UV system is not initiated entirely by default, and that the U-specific region is required to fully express the female developmental program. Taken together, our results provide the first illustration of how male- and female-specific developmental programs in a haploid sex determination system may, at least partially, be uncoupled from sex chromosome identity.

## Results

### Identification of a genetically male line exhibiting female-like morphology

In the giant kelp *M. pyrifera*, sex is determined genetically during the (haploid) gametophyte generation. Meiosis occurs during the sporophyte generation, producing haploid spores that develop into either male or female gametophytes. Male and female gametophytes exhibit marked sexual dimorphism in terms of size of vegetative cells and reproductive structures (Westermeier et al. 2007; Lipinska, Ahmed, et al. 2015; Luthringer et al. 2015). Male gametophytes produce small male gametes (sperm) in specialised sexual structures called antheridia, whereas female gametophytes produce large female gametes (eggs) in oogonia (Westermeier et al. 2007).

During a screen of colchicine treated *M. pyrifera* gametophytes, we identified a genetically male line (Mpyr-13-4) that exhibited morphological features resembling those of female gametophytes (Figure 1; Figure S1). To determine whether the observed phenotypes were due to polyploidy or aneuploidy induced by the colchicine treatment, we measured the ploidy level of Mpyr-13-4 using flow cytometry. This analysis showed no evidence for chromosome doubling (Figure S2), indicating that the colchicine treatment had not caused large scale chromosomic modifications.

**Figure 1.**
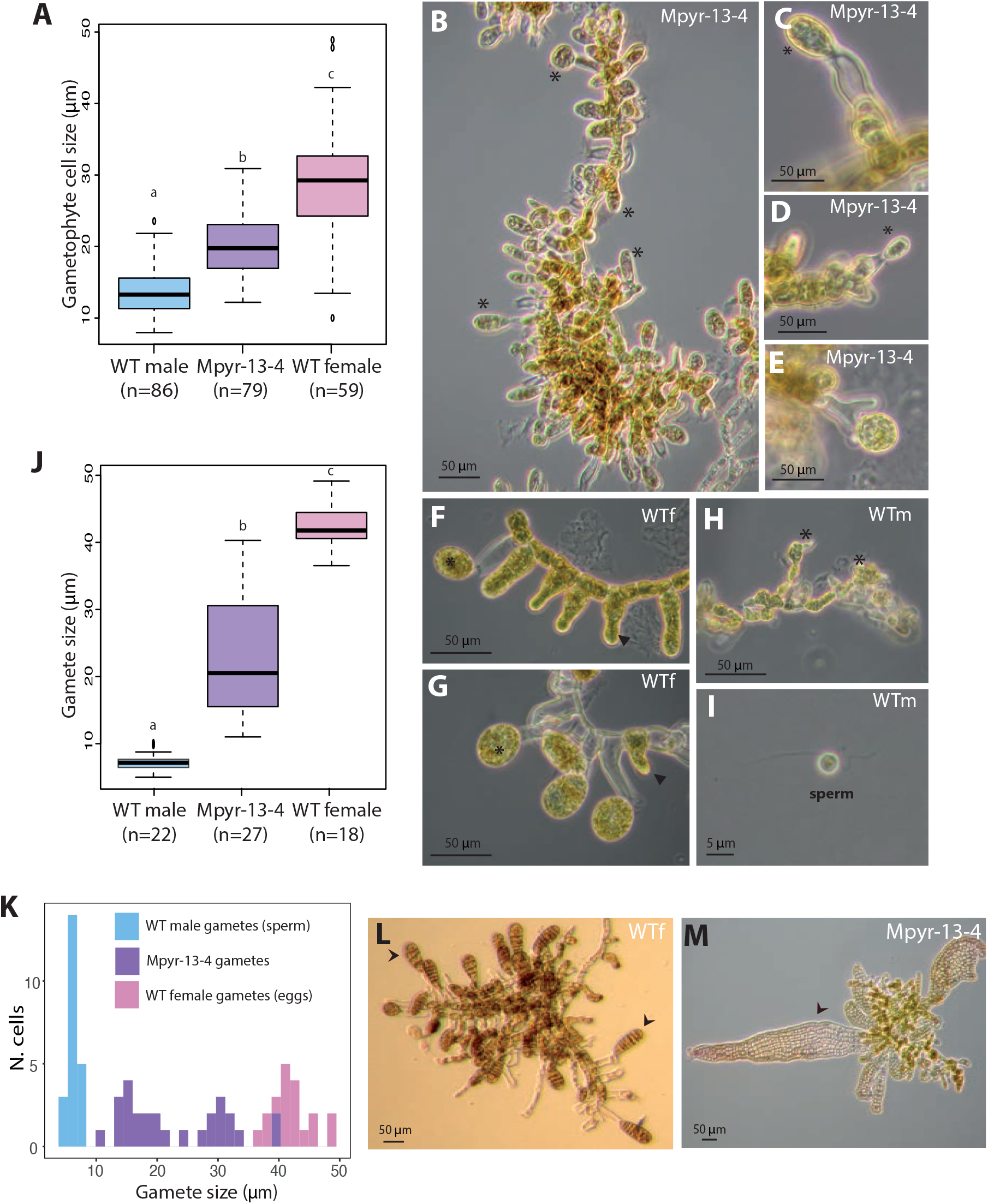
Phenotypic characterisation of wild-type male and female gametophytes and the variant Mpyr-13-4 line. (A) Sizes of gametophyte cells in wild-type male, wild-type female and Mpyr-13-4 individuals. (B-E) Fertile Mpyr-13-4 gametophytes showing several egg-like structures of different sizes (asterisks). (F-G) Fertile wild-type female gametophyte with extruded eggs (asterisks) and oogonia (arrow head); (H) Fertile wild-type male gametophyte with antheridia (asterisks) and sperm (I). (J) Gamete sizes in wild-type males, wild-type females and Mpyr-13-4. Different letters above the plots indicate significant differences (Wilcoxon rank sum test, p-value<0.001). (K) Distribution of the gamete sizes for wild-type male, wild-type female and Mpyr-13-4 variant lines. (L) Parthenogenetic sporophytes in wild-type females (arrow heads). (M) Parthenogenetic sporophytes in Mpyr-13-4 (arrow heads).

### Phenotypic characterisation of the feminized line compared with wild-type males and females

A detailed morphometric study of gametophytes of the Mpyr-13-4 strain showed the cells of this strain were significantly larger than those of the wild-type male strain (Wilcox test W = 31, p-value = 6.27e-10) (Figure 1A), exhibiting intermediate size between those of cells of male and female wild-type strains.

Following induction of gametogenesis, Mpyr-13-4 gametophytes formed reproductive structures (Figure 1B-1E) that strongly resembled wild-type female oogonia (Figure 1F-1G) and not male antheridia (Figure 1H-1I). At maturity, Mpyr-13-4 gametophytes produced egg-like cells, lacking the flagella typically present in male gametes, with diameters ranging from 10-40 µm. Wild-type eggs are typically around 40 µm in diameter (Figure 1K).

In absence of fertilisation by gametes of the opposite sex, wild-type female gametes of *M. pyrifera* initiate parthenogenetic development within 48 hours (Figure 1L). Wild-type male gametes do not undergo parthenogenesis. Mpyr-13-4 gametes, regardless of their size, initiated parthenogenetic development within 48 hours (Figure 1M).

Note that the female-like phenotypes of Mpyr-13-4 were stably maintained through (asexual) vegetative reproduction, and were not affected by culture conditions (10°C, 15°C, high/low white light, red light).

Taken together, these analyses indicated that despite being genetically male, Mpyr-13-4 individuals exhibit a feminised phenotype, with several phenotypic features more typical of females, including gametophyte cell size, presence of oogonia-like structures, gamete cell size and parthenogenetic capacity.

### Mpyr-13-4 gametes do not produce the male-attracting pheromone lamoxirene

In order to test if gametes produced by the feminised strain were fully functional female gametes, we performed test crosses (Westermeier et al. 2007). When Mpyr-13-4 gametes were confronted with active sperm from an unrelated, wild-type *M. pyrifera* male line, no gamete interaction was observed, and no zygotes were produced.

Wild-type female gametes attract male gametes by producing a pheromone (lamoxirene; (Maier et al. 2001). This pheromone is responsible both for the release of sperm from male antheridea and for the attraction of sperm towards the eggs. We therefore investigated whether Mpyr-13-4 gametes were capable of producing lamoxirene. MS/MS analysis of *M. pyrifera* detected lamoxirene in control wild-type female gametophytes, but failed to detect lamoxirene or other C11 hydrocarbons in the feminised Mpyr-13-4 fertile gametophytes (Figure 2). This observation suggests that despite presenting phenotypic features typical of a female, the Mpyr-13-4 strain is not fully recapitulating the female developmental program.

**Figure 2.**
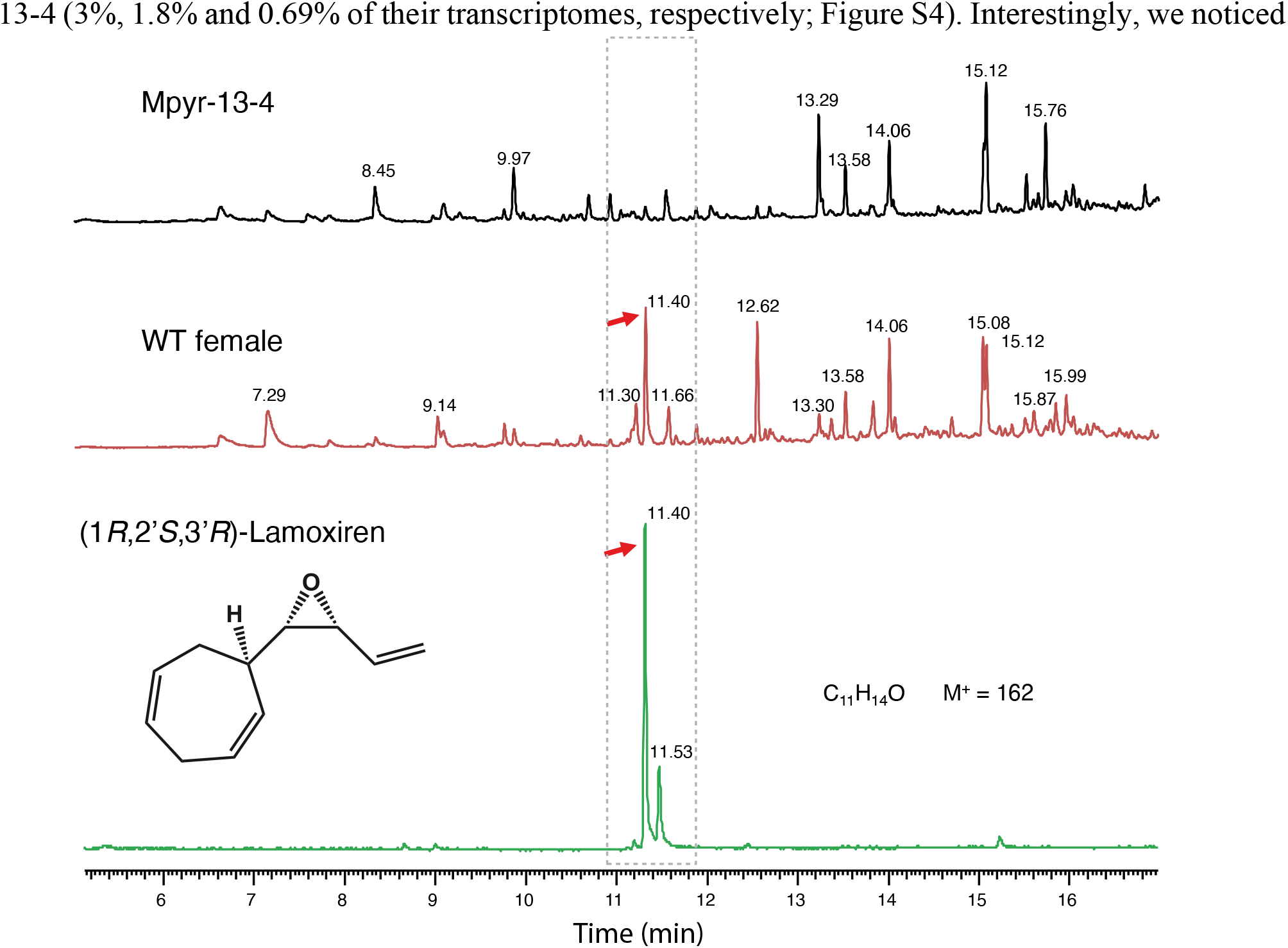
GS/MS assay for lamoxirene detection in wild-type females and Mpyr-13-4 lines. Red arrows indicate the peak of lamoxirene

### Gene expression in Mpyr-13-4 compared with wild-type male and female gametophytes

To determine whether the morphological feminisation of the variant male strain was associated with changes in terms of transcriptional landscape compared with wild-type male and female gametophytes, transcript abundances were measured by RNA-seq analysis of wild-type male, wild-type female and Mpyr-13-4 gametophytes (Table S1 and methods for details).

This analysis identified 17,922, 18,436 and 17,744 expressed genes (defined as TPM>5^th^ percentile) in males, females and the Mpyr-13-4 strain, respectively, of the total of 22,242 genes that have been annotated in the *M. pyrifera* genome (Lipinska et al. 2019); Table S2, Figure S3, S4). Hence, 81%, 82% and 80% of the total number of annotated genes were detected as being expressed in wild-type male, wild-type female and Mpyr-13-3 gametophytes, respectively.

The majority (app. 94%) of the transcriptome was expressed in all three samples but a slightly greater number of genes were uniquely expressed in wild-type females than in wild-type males and Mpyr-13-4 (3%, 1.8% and 0.69% of their transcriptomes, respectively; Figure S4). Interestingly, we noticed that the variant Mpyr13-4 strain shared more expressed genes with wild-type females than with wild-type males (610 and 327 genes respectively; Figure S4). Moreover, a sample distance matrix showed a closer relationship between wild-type female and Mpyr-13-4 samples compared with wild-type male (Figure S5).

### Sex-biased gene expression in wild-type and Mpyr-13-4 lines

In order to examine if the line Mpyr-13-4 was feminized and/or demasculinized at the level of gene expression, we next analysed the patterns of expression of sex-biased genes in each of the samples.

DEseq2 analyses identified 5442 genes that were differentially expressed between males and females, indicating that a considerable proportion (24.5%) of the transcriptome of *M. pyrifera* exhibits sex-biased expression (Table S2, S3). Approximately the same numbers of genes were found to be male-biased (2785) and female-biased (2657) (Table S2, S3, Figure 3A). Consistent with the female-like phenotype of Mpyr13-4, more differentially expressed genes were identified when this strain was compared with wild-type males (20.8%) than when it was compared with wild-type females (5.8%) (Table S2, S3, Figure 3A).

**Figure 3.**
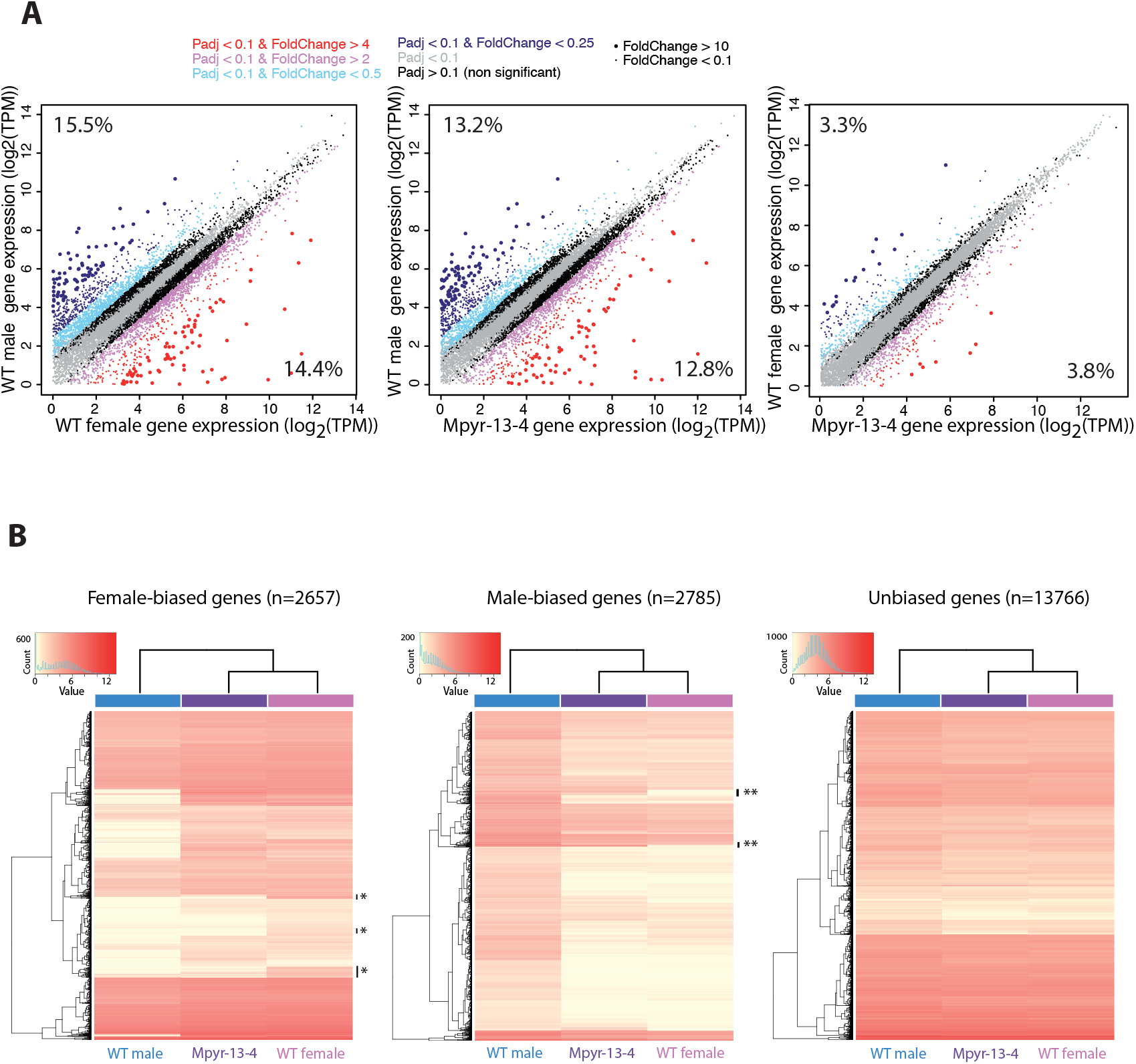
Sex-biased gene expression (A) Comparisons of gene transcript abundances (log2TPMs) in wild-type male, wild-type female and the Mpyr-13-4 line. Genes whose expression was 0 in one of the samples were removed from the plot. The percentages of biased genes in each sample are indicated in the upper left and lower right-hand corners. (B) Heat maps and hierarchical clustering of gene expression for wild-type male, wild-type female and Mpyr-13-4 variant line. Shown is the relative expression for autosomal male-biased, female-biased and unbiased genes. Hierarchical gene clustering is based on Euclidean distance for average log2 expression of each gene for the three samples.

Hierarchical clustering of expression levels was used to visualize gene transcription patterns for wild-type male, female and variant Mpyr-13-4 samples. Wild-type female and Mpyr-13-4 samples clustered together both when sex-biased gene expression was analysed and when the expression of non-sex-biased genes (unbiased genes) was analysed, indicating that the transcriptome of the Mpyr-13-4 strain was more similar to that of a wild-type female than to that of a male (Figure 3B). Principal component analysis (PCA) further supported this conclusion (Figure S3).

In order to examine how sex-biased gene expression is affected by the phenotypic feminisation of the variant Mpyr-13-4 strain, we next focused on the sets of genes that had been defined as sex-biased in the wild-type strains. The median level of expression (measured as log_2_(TPM+1)) of the male-biased gene set in Mpyr-13-4 gametophytes was 79.4% lower than that observed in wild-type males; (Wilcoxon test, p<2.2E-16, Figure 4A), suggesting these genes are transcriptionally “de-masculinized” in Mpyr-13-4. In contrast, female-biased genes were expressed at a higher level in Myr-13-4 (31.6% more) than in wild-type males (Wilcoxon test, p<2.2E-16, Figure 4A), suggesting that Mpyr-13-4 is transcriptionally “feminised”. Finally, genes with unbiased expression (n=13766) showed no average change in expression between wild-type male and females and variant strain (Figure S6).

**Figure 4.**
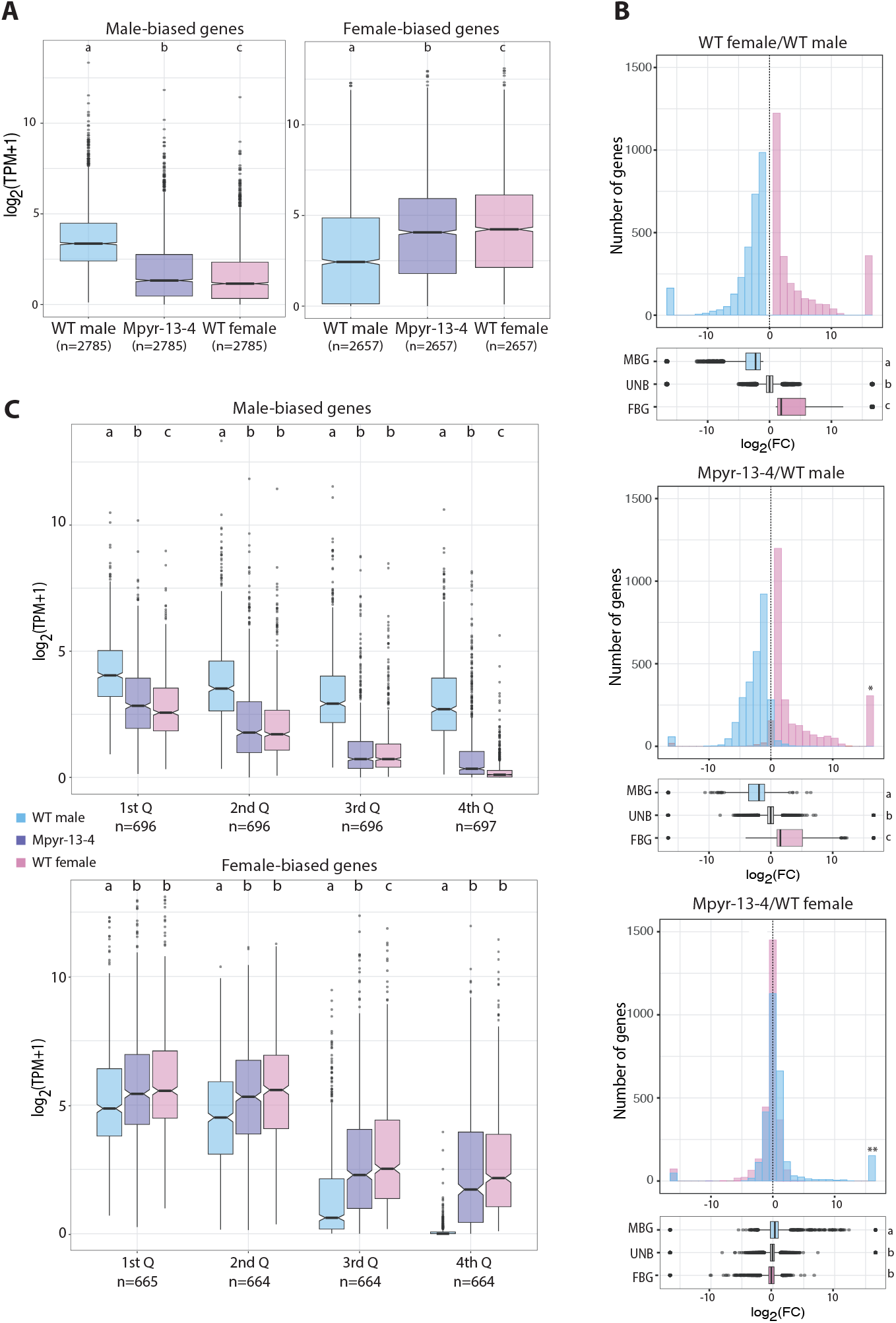
(A) Abundances of transcripts (TPM) of male- and female-biased genes in wild-type males, wild-type females and Mpyr-13-4. (B) Transcript abundance fold changes in pairwise comparisons between wild-type males, wild-type ¬¬females and in Mpyr-13-4. Upper panel: wild-type female versus wild-type male. Positive values in the x axis correspond to higher expression in females, negative values to higher expression in males. Middle panel: gene expression in Mp-13-4 versus wild-type male gametophytes. Positive values on the x axis correspond to higher expression in the variant Mpyr-13-4, negative values to higher expression in wild-type males. Lower panel: Positive values on the x axis correspond to higher expression in Mpyr-13-4, negative values to higher expression in wild-type females. Colours indicate male-biased (blue), female-biased (pink) and unbiased (grey) genes. For clarity, unbiased genes are omitted from the histogram. Different letters indicate significant differences (Wilcoxon rank sum test, p-value < 0.01). (C) Male-biased (upper panel) and female-biased (lower panel) genes ranked by level of bias (log_2_TPM+1). Numbers of genes are indicated below the plots. Note that only genes with TPM>0 are included in the plots. Significant differences detected following Wilcoxon tests (p-values < 0.01) are indicated as different letters above the plots.

Transcript abundance fold changes in pairwise comparisons between wild-type males, wild-type females and in Mpyr-13-4 also revealed a pattern of simultaneous feminisation and de-masculinisation of the transcriptome of Mpyr-13-4 (Figure 4B).

However, we noticed that a subset of female-biased genes that was completely silenced in wild-type males was also not expressed in Mpyr-13-4 (Figure 3B, Figure 4B). Conversely, male biased genes that were silenced or lowly expressed in wild-type females were expressed in Mpyr-13-4 (Figure 3B, Figure 4B).

Taken together, these observations indicate that although the transcriptome of Mpyr-13-4 was clearly feminised and de-masculinised, there was not a complete shift to the female transcriptional program.

To examine in more detail the relationship between degree of sex-biased gene expression (fold change in transcript abundance between the wild-type female and male) and transcript abundance, the sex-biased genes were grouped according to fold-change (FC) differences between the male and female samples and median transcript abundances plotted for wild-type males, wild-type females and Mpyr-13-4. The global pattern of de-masculinisation and feminisation of Mpyr-13-4 transcription appeared to be strongly correlated with the degree of sex-bias (Figure 4C). Down-regulation of the male-biased gene set in Mpyr-13-4 was more pronounced for the genes that showed the highest levels of male-bias, suggesting that the male-biased genes that exhibit the greatest fold changes make the greatest contribution to male-specific traits in the variant strain. Similarly, stronger upregulation was observed in Mpyr-13-4 for the genes that exhibited the highest levels of female-bias (Figure 4C).

Taken together, these results indicate the transcriptome of Mpyr-13-4 is both de-masculinised and feminised, with the strongest effect on genes that exhibit strong sex-biased expression patterns (i. e. highest FC between wild-type male and female).

### Predicted functions of genes involved in feminisation

An analysis of gene ontology (GO) terms associated with the sex-biased genes and with the genes differentially expressed in Mpyr-13-4 was carried out using BLAST2GO (C(Conesa and Götz 2008). The aim of this analysis was to search for enrichment in particular functional groups and to relate gene function to phenotypic feminisation. Significant enrichment of specific GO categories related to microtubule- and flagella-related processes was detected for genes that were down-regulated in Mpyr-13-4 compared with wild-type males (Table S5). These genes may be involved in the production of flagellated gametes, which are not produced by Mpyr-13-4. Note that the these two GO categories were also enriched in the set of male-biased genes expressed in male gametes identified by (Lipinska et al. 2013) and in fertile male gametophytes (Lipinska, Cormier, et al. 2015; Lipinska et al. 2019). The female-biased genes that were upregulated in Mpyr-13-4 compared to wild-type males were enriched in GO terms related to metabolism and photosynthesis (Table S5).

We identified a set of 312 genes likely involved in the female developmental program whose regulation depends on the presence of the U-specific region (genes that were over-expressed in wild-type females compared with wild-type males and Mpyr-13-4). GO term analysis revealed an enrichment in metabolic pathways-related and oxidation-reduction functions (Table S5).

### Evolutionary analysis of genes involved in the Mpyr-13-4 phenotype

Analyses of the evolutionary dynamics of sex-biased genes in other organisms have shown that they evolve more rapidly than unbiased genes, which is expected because sexually dimorphic traits contribute to reproductive success (reviewed in (Parsch and Ellegren 2013a). We therefore tested if the genes involved in sexual differentiation in the giant kelp and in particular the genes involved in the feminisation and de-masculinisation phenotype exhibited by Mpyr-13-4 showed evidence for different evolutionary rates compared with unbiased genes.

To test for differences in rates of evolutionary divergence between different categories of sex-biased and unbiased genes, we calculated levels of nonsynonymous (dN) and synonymous (dS) substitution using pairwise comparisons with orthologs from another kelp species, *Saccharina japonica*. The results of this analysis indicated that genes exhibiting male-biased expression patterns in the giant kelp gametophytes evolve significantly faster (i.e., had higher dN/dS values) than female-biased or unbiased genes (Mann–Whitney U test, P < 0.01) (Figure 5). Female-biased genes were found to have evolved more slowly than unbiased genes, and this was due to a combination of lower dN and higher dS than unbiased genes. Sex-biased genes that were differentially expressed in Mpyr-13-4 compared with wild-type males or wild-type females exhibit evolutionary patterns that were not significantly different from male- and female-biased genes, respectively (Figure S7).

**Figure 5.**
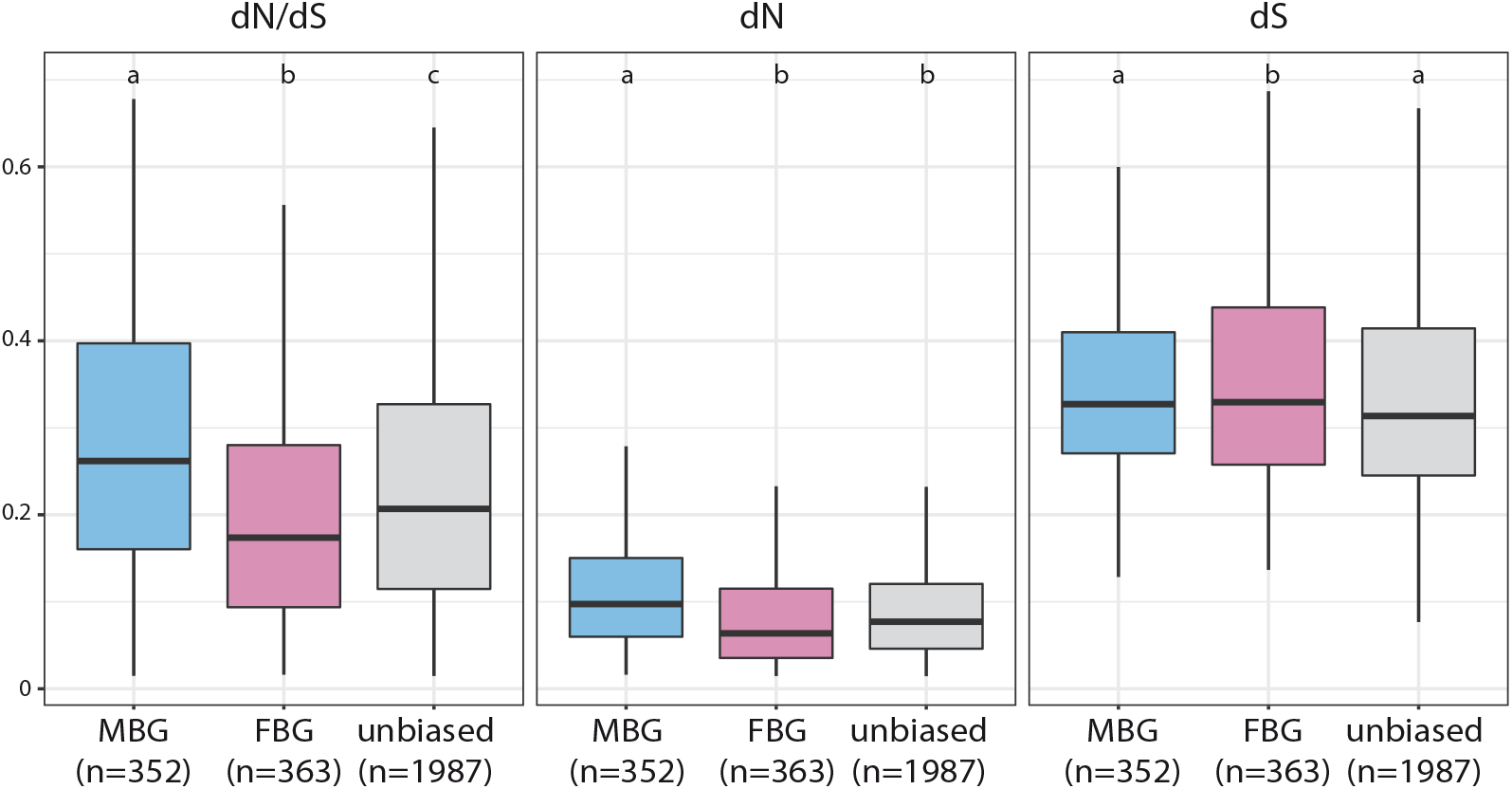
Rates of evolution of male-biased, female-biased and unbiased genes. Pairwise dN, dS and dN/dS ratios were calculated by comparing orthologous sequences from M. pyrifera and S. japonica (Ye et al., 2015). From left to right, ratio of nonsynonymous to synonymous substitutions (dN/dS), synonyms substitutions (dS) and nonsynonymous substitutions (dN). Whiskers extend to the most extreme data point, excluding outliers that exceeded 1.5x the interquartile range. Significant differences detected following Wilcoxon tests (p-values < 0.01) are indicated as different letters above the plots.

Highly expressed genes are often observed to exhibit lower dN/dS values (Drummond et al. 2005; Cherry 2010; Slotte et al. 2011) and overall, female-biased genes were more highly expressed than unbiased or male-biased genes (pairwise Wilcoxon test p-value < 2.2e-16; Figure S6). Female-biased genes are notorious for exhibiting slower evolutionary rates compared with male biased genes, and this has been related to the fact they are more pleiotropic (Grath and Parsch 2016). No data is available for different tissues or several life cycle stages of the giant kelp so we could not measure breadth of expression as a proxy for pleiotropy (Uebbing et al. 2016; Lipinska et al. 2019) however, it is interesting to note that female-biased genes are expressed both in gametophyte and in sporophyte stages whereas male biased genes appear to have a more restricted expression to male gametophytes (Figure S8).

### Expression of male SDR genes in the Mpyr-13-4 line

The feminised phenotype of the Mpyr-13-4 strain could be consistent with modification of the expression of a sex-determining gene or genes carried by the V-specific region. We thus specifically focused on the expression patterns of previously identified male-linked *M. pyrifera* genes (Lipinska et al. 2017). For the majority (8 out of 10) of these genes, similar transcript abundances were detected in wild-type males and the Mpyr-13-4 strain (Figure 6). However, two genes, gHMG.13001750 and gSDR.13001840, were markedly downregulated in the Mpyr-13-4 line compared with wild-type males. The orthologs of these two genes in *Ectocarpus* are Ec-13-001750 and Ec-13-001840, respectively. Interestingly, Ec-13-001750 is the candidate master-sex-determining gene in *Ectocarpus* sp. (Ahmed et al. 2014).

**Figure 6.**
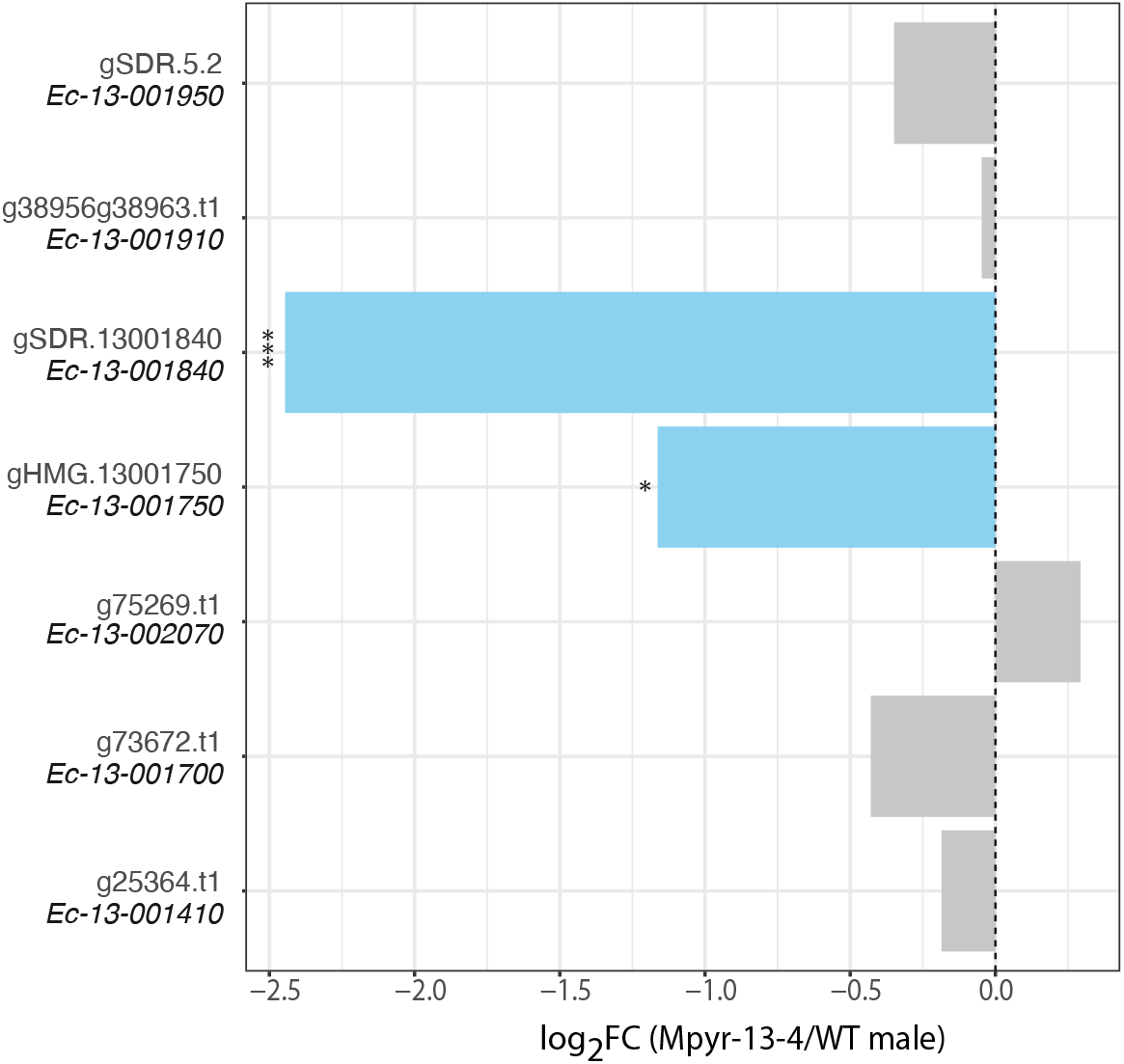
Differential expression (DESeq2) of M. pyrifera orthologues of Ectocarpus male SDR genes in Mpyr-13-4 versus wild-type males. Negative values represent downregulation (log2FC) in the variant Mpyr-13-4 strain relative to the wild-type male.

## Discussion

### Transcriptome de-masculinisation and feminisation underlie the phenotype of Mpyr-13-4

Gametophytes of the giant kelp variant line Mpyr-13-4 exhibited female phenotypic characteristics despite being genetically male. We used wild-type male and female lines to examine whether the degree of sex-biased gene expression is associated with phenotypic sexual dimorphism, and to understand the role of gene expression in encoding the variant morphology. Our results reveal a simultaneous de-masculinisation and feminisation of the transcriptome of the Mpyr-13-4 line and underline the association between sex-specific phenotypes and sex-specific transcriptomic patterns. The overall pattern of de-masculinisation and feminisation of Mpyr-13-4 transcription was strongly correlated with the degree of sex-bias (measured as fold-change in expression between wild-type male and wild-type female). De-masculinisation of gene expression in the variant line was more pronounced for genes that exhibited the strongest male-bias (highest fold-change), possibly suggesting that the most extreme male-biased genes make the greatest contribution to male-specific traits in the giant kelp. Similarly, feminisation of the variant transcriptome manifested itself as increased expression of female-biased genes, and was more marked for the genes with highest fold changes between wild-type male and wild type female. A similar tendency was observed in animals with ZW systems, where masculinisation of gene expression was associated with increased male sexual dimorphism, and the genes with most extreme sex-bias make the greatest contribution to sex-specific traits (Pointer et al. 2013). Our results therefore extend the link between sexually dimorphic transcription and sexually dimorphic phenotypes to haploid sex determining systems.

Despite a clear de-masculinisation and feminisation of the transcriptome of the variant line, there was not a full shift to the female transcriptomic program and, accordingly, the phenotype of the variant line did not fully recapitulate the female developmental program.

### The female SDR may be required to fully express the female developmental program

Despite the phenotypic resemblance to a female, the Mpyr-13-4 gametophyte produced non-functional eggs, which failed to attract male gametes. MS/MS analyses revealed absence of pheromone production in the eggs-like cells of the variant line, so it appears that the factor(s) required to initiate the pheromone pathway is absent or not functional in this line. Pheromone production may be regulated directly (or indirectly) by a gene(s) located in the SDR, similar to the case of yeast where the mating type linked Ste-20 regulates mating through pheromone response pathways (Leberer et al. 1992). One gene within the *M. pyrifera* sex-linked region is a Ste-20-like serine-threonine kinase (Lipinska et al. 2017), but currently there is no evidence for a role of this gene in pheromone production in brown algae.

Interestingly, examination of the predicted functions of the list of genes that were over-expressed in wild-type females but not in wild-type males nor Mpyr-13-4, i. e. presumably the genes that may be regulated by the U-specific region, revealed an enrichment in metabolism-related and oxidation-reduction functions. In pheromone producing stages of yeast, significant increases in the transcript levels of a multitude of metabolic enzymes are also observed (Williams et al. 2016). It is therefore possible that among these wild-type female-limited genes there are components of the giant kelp pheromone biosynthetic cascade or its regulators.

Taken together, our observations are coherent with incomplete sex-reversal of Mpyr-13-4, and suggest that the female SDR is required to fully express the female program of development in this UV system.

### Role of the SDR and autosomal gene expression in the initiation of sex-specific developmental programs

Although Mpyr-13-4 is not a fully male-to-female sex reversed line, analysis of the Mpyr-13-4 line contributed to understanding the role of sex-linked genes and autosomal gene expression in the initiation of the male and female developmental programs. Our observations indicate that many of the female developmental features depend exclusively on autosomal gene expression and do not require the presence of the female SDR. One particularly interesting feature was the capacity of the variant line to undergo parthenogenesis, a trait that is typical of females of oogamous brown algal species (Luthringer et al. 2015). In the model brown alga *Ectocarpus*, parthenogenesis is a complex genetic trait under the control of the SDR, together with at least two additional autosomal loci (Mignerot et al. 2019). It has been suggested that either the male SDR produces a repressor of parthenogenesis, or, alternatively, the female SDR produces an activator of parthenogenesis (in either case the activator or repressor could be directly encoded by the SDR or produced indirectly as part of the male or female sex-differentiation programs). Taking into consideration the parthenogenetic development of the egg-like cells of the Mpyr-13-4 line (which lacks the female SDR), observed in this study, it appears clear that the female SDR is not required for parthenogenesis to be initiated in kelps. Therefore, parthenogenesis may involve a repressor being produced by the male SDR, as part of the male-differentiation program or, alternatively, parthenogenesis could be induced as part of the female sex-differentiation program but independently of the female SDR.

### The transcriptional and evolutionary landscape underlying sexual differentiation in the giant kelp

Sex-biased gene expression has been characterised for the brown alga *Ectocarpus*, a near-isogamous species with a low level of sexual dimorphism (Lipinska, Cormier, et al. 2015; Luthringer et al. 2015). Here we show that substantially more genes are sex-biased in the giant kelp (about 24% of the transcriptome, compared with less than 10% in *Ectocarpus* (Lipinska, Cormier, et al. 2015). The higher proportion of sex-biased genes in the giant kelp is consistent with the higher level of phenotypic sexual dimorphism in this organism, where male and female gametophytes have clearly distinct morphologies (Muller et al. 1979).

The rapid rates of evolution sex-biased genes observed in animals and plants are thought to be due to a combination of natural selection, sexual selection, and relaxed purifying selection (Parsch and Ellegren 2013b; Grath and Parsch 2016). Our study revealed that the genes involved in sex-specific development in the giant kelp show a broadly similar pattern of sequence evolution to that seen in animals, with male-biased genes showing elevated rates of evolution compared to female-biased and unbiased genes, consistent with the notion that sex-specific selection may be stronger in males than females in this species. This is contrasts with previous studies in *Ectocarpus* sp., where both male- and female-biased genes exhibited similar and faster evolutionary rates than unbiased genes (Lipinska, Cormier, et al. 2015). Kelps have more conspicuous sexual dimorphism with males producing higher amount of sperm (Muller et al. 1979) compared with eggs, providing scope for sperm competition and higher levels of sexual selection specifically in males. Moreover, in *M. pyrifera*, as in other kelp species, female gametes remain attached to the parental gametophyte and male gametes swim towards the eggs, attracted by the pheromone. Therefore, males have to produce large quantities of rapidly swimming male gametes, and there may be more scope for male competition than for female competition.

### V-linked genes potentially required for male developmental patterns

Two genes located within the giant kelp male SDR exhibited a significant reduction in transcript abundance in the Mpyr-13-4 line, suggesting their involvement in the variant phenotype. One of these genes is a HMG-domain protein coding gene, with orthologs present with the VSR across a range of brown algae and thought to be the master male sex determining gene for this group of organisms (Ahmed et al. 2014; Lipinska et al. 2017). The other gene is a membrane-located protein-coding gene with a putative sugar binding and cell adhesion domains. Both genes belong to a small set of genes that have been conservatively sex linked across all the brown algae studied so far (Lipinska et al. 2019). Our observations suggest that these ancestrally male-linked genes are involved in the development of male functional characteristic, although mechanistic studies will be necessary to fully validate their role in male sex determination and differentiation.

## Materials and Methods

### Biological material

A *Macrocystis pyrifera* sporophyte was collected at Curanoe, Chile. Meiospores were isolated and clonal cultures of male and female gametophytes were established and propagated as described by (Westermeier et al. 2007). Male and female gametophyte clones (Westermeier et al. 2010) were selected for the present study (Figure S1). Axenic sub-clones were initiated by antibiotic treatment as described by (Müller et al. 2008) and maintained on 1% agar in seawater with transfers every 3-months. For clone isolation, culture medium was prepared with a commercial salt mixture (hw-Professional, Wiegandt, Krefeld Germany) in demineralized water and adjusted to 3% salinity with an optical refractometer. Nutrients were added with 20 ml/L PES-enrichment (Starr and Zeikus 2004; Coelho et al. 2012). Colchicine treatment was performed using a disk of filter paper of 6 mm diameter loaded with 1 mg of colchicine (Fluka, Honeywell Research Chemicals), which was placed in the centre of an agar plate filled with gametophyte material, with good contact between agar and paper in order to allow diffusion of the colchicine into the agar. The agar plates were sealed with parafilm and subjected to culture conditions (12 ± 2°C and 2-3 µE m^-2^ sec^-1^ from daylight type fluorescent lamps for 14:10 (light:dark) cycles for 12 to 16 weeks until selected regenerates were isolated.

### Pheromone measurements

Female eggs of *M. pyrifera* produce a pheromone (lamoxirenee) that attracts male gametes. To induce gametogenesis in wild-type female and Mpyr-13-4 lines, light intensity was increased to 30 µE m^-2^ sec^-1^ and culture medium was refreshed every 4 days. Oogonia and eggs were produced between 9-16 days after culture in these conditions.

For each of the tested strains, 25 mL of fertile gametophyte cultures containing 6.5 × 10^4^ eggs (wild-type female) or 3.1 × 10^5^ eggs (Mpyr-13-4) were introduced in 50 mL Greiner Cellstar tissue culture tubes placed horizontally to increase surface area.

All algal pheromones known are hydrophobic, cycloaliphatic unsaturated hydrocarbons, comprising eight to eleven carbon atoms [1]. The sperm-releasing pheromone in the Laminariales carries an additional epoxy moiety, but is still a volatile and hydrophobic compound (Maier et al. 2001). To efficiently trap these pheromones, which are released in only minute amounts from fertile gametes, Solid Phase Micro-extraction (SPME) was used (Maier et al. 1996). SPME fibers and the holder were obtained from Supelco (Bellefonte, PA, USA). For volatile extraction, a poly-dimethyl-siloxane (100 µm PDMS) fiber was used (red fiber). Prior to use, the SPME fibers were conditioned according to the manufacturer’s instructions. After 14 h exposure time of the fiber in the culture medium, the fiber was thermally desorbed in the gas chromatographer (GC/MS) injection port followed by separation of the volatiles under programmed conditions using an ISQ LT and Trace 1310 (Thermo Fisher Scientific GmbH, Dreieich, Germany) device equipped with a ZB5 column (30 m, 0.25 mm I.D., 0.25 µm film thickness) linked to a guard column (10 m, Phenomenex, Aschaffenburg, Germany). Helium (1.5 ml.min^-1^) served as the carrier gas. Separation of compounds was achieved under programmed conditions from 50°C (2 min isotherm), followed by heating at 10°C min^-1^ to 200°C and at 50 °C min^-1^ to 280°C. The GC injector (splitless, splitless time 2 min), transfer line and ion source were set at 230, 280 and 250°C, respectively. Mass spectra were recorded in electron impact (EI) mode at 70 eV, 35–350 m/z.

### Generation of transcriptomic sequence data

The algal strains used, sequencing statistics, and accession numbers are listed in Table S1.

RNA-seq analysis was carried out to compare the relative abundances of gene transcripts in the different samples. For each sample, total RNA was extracted from 2 independent bulks of app. 1000 male individuals and 2 bulks of 1000 female individuals (two biological replicates for each sex) using the Qiagen Mini kit (http://www.qiagen.com) as previously described (Lipinska, Cormier, et al. 2015; Arun et al. 2019). RNA from wild-type male and female and variant Mpyr-13-4 line pools was extracted using the protocol described by (Apt et al. 1995). RNA quality and quantity were assessed using an Agilent 2100 bioanalyzer, associated with an RNA 6000 Nano kit.

For each replicate, the RNA was quantified and cDNA was synthesized using an oligo-dT primer. The cDNA was fragmented, cloned, and sequenced by Fasteris (CH-1228 Plan-les-Ouates, Switzerland) using an Illumina Hi-seq 2000 set to generate 150-bp single-end reads. Data quality was assessed using the FastQC (Wingett et al. 2018). Reads were trimmed and filtered using Trimmomatic (B(Bolger et al. 2014) with average quality > 28, a quality threshold of 24 (base calling) and a minimal size of 60 bp.

Filtered reads were mapped to the *M. pyrifera* genome (Lipinska et al. 2019) using TopHat2 (Kim et al. 2013) with the Bowtie2 aligner (Langmead and Salzberg 2012).

More than 70% of the sequencing reads for each library could be mapped to the genome (Table S1). The mapped sequencing data were then processed with FeatureCounts (Liao et al. 2014) to obtain counts for sequencing reads mapped to exons and counts by gene. Transcript abundances, measured as transcript per million (TPM) were strongly correlated between biological replicates of each sample (Figure S5).

Expression values were represented as transcript per million (TPM); any genes with TPM value above the 5^th^ percentile is considered as expressed. This resulted in a total of 19208 genes with express in at least one of the three strains.

### Identification of sex-biased genes

The filtering steps described above yielded a set of expressed genes in the transcriptome that were then classified based on their sex-expression patterns. Differential expression analysis was performed with the DESeq2 package (Love et al. 2014) (Bioconductor). Genes were considered to be male-biased or female-biased if they exhibited at least a twofold difference in expression between generations with a false discovery rate (FDR) of < 0.05. Sex-biased genes were defined as sex-specific when the TPM was below the fifth percentile for one of the sexes. Full lists of sex-biased genes can be found in Table S3.

### Evolutionary analysis

To estimate evolutionary rates (non-synonymous to synonymous substitutions, dN/dS) we used single copy ortholog genes between *Macrocystis pyrifera* and *Saccharina japonica* described in (Lipinska et al. 2019). Protein sequences were aligned with Tcoffee (M-Coffee mode) (Notredame et al. 2000) and translated back to nucleotide sequence using Pal2Nal (Suyama et al. 2006). Gapless alignments that exceeded 100 bp were analysed with CodeML (F3×4 model of codon frequencies, runmode = −2) implemented in the Phylogenetic Analysis by Maximum Likelihood (PAML4) suit (Yang 2007)). Genes with saturated synonymous substitution values (dS > 1) were excluded from the analysis.

### Flow cytometry

Flow cytometry was performed as described before for brown algal tissues (Bothwell et al. 2010). Gametophyte tissue was finely cut with a razor blade and nuclei were isolated by suspension in nuclei buffer (30 mM MgCl, 120 mM trisodium citrate, 120 mM sorbitol, 55 mM 4-(2-hydroxyethyl) piperazine-1-ethanesulfonic acid (HEPES), pH 8, 5 mM EDTA supplemented with 0.1% (v/v) Triton X-100 and 5 mM sodium bisulfite; pH 8.0), and their DNA content was measured immediately by flow cytometry. Between 600 and 13, 200 nuclei were analysed in each sample. Wild-type gametophytes were considered to be haploid and were used as an internal reference for the determination of ploidy. The nucleic acid-specific stain SYBR Green I (http://www.invitrogen.com) was used at a final dilution of 1:10,000. Samples were analysed using a FACSort flow cytometer (http://www.bsbiosciences.com).

## Supplemental Figures

**Figure S1.**
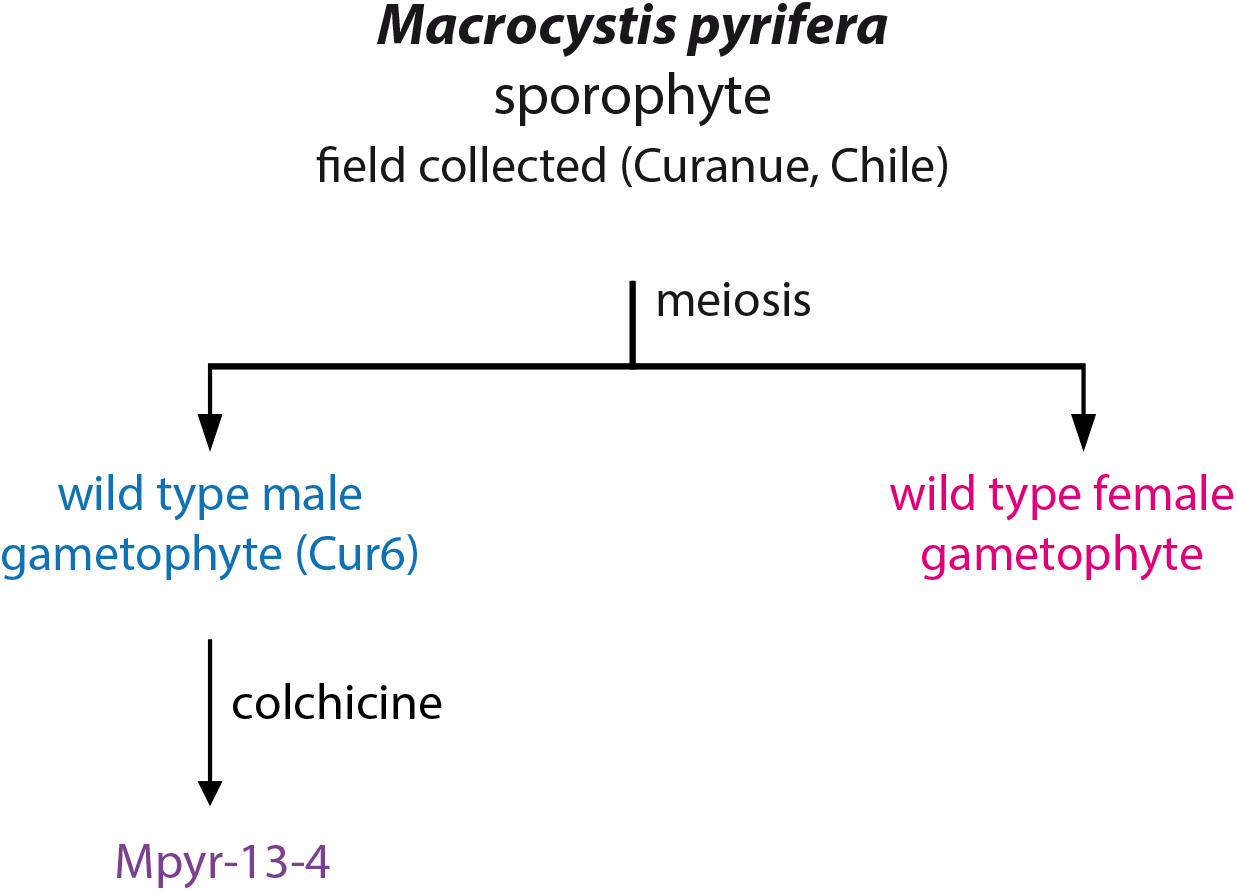
Pedigree of the strains used in this study.

**Figure S2.**
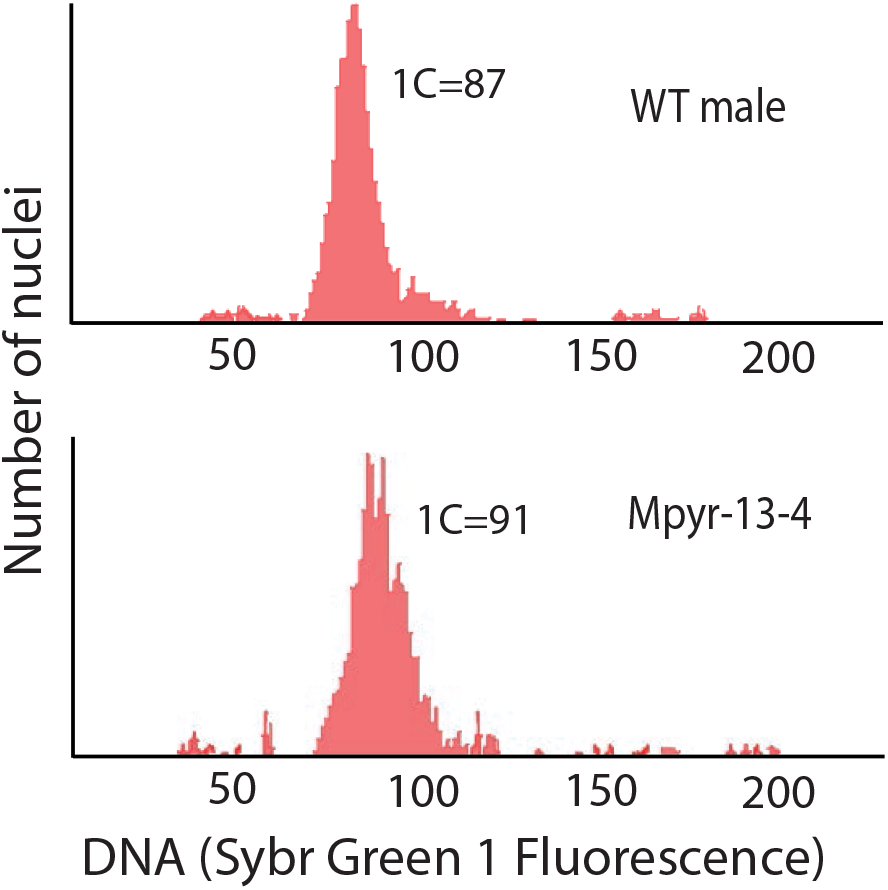
Flow cytometry analysis of a wild-type male line (top) and the Mpyr-13-4 line (bottom).

**Figure S3.**
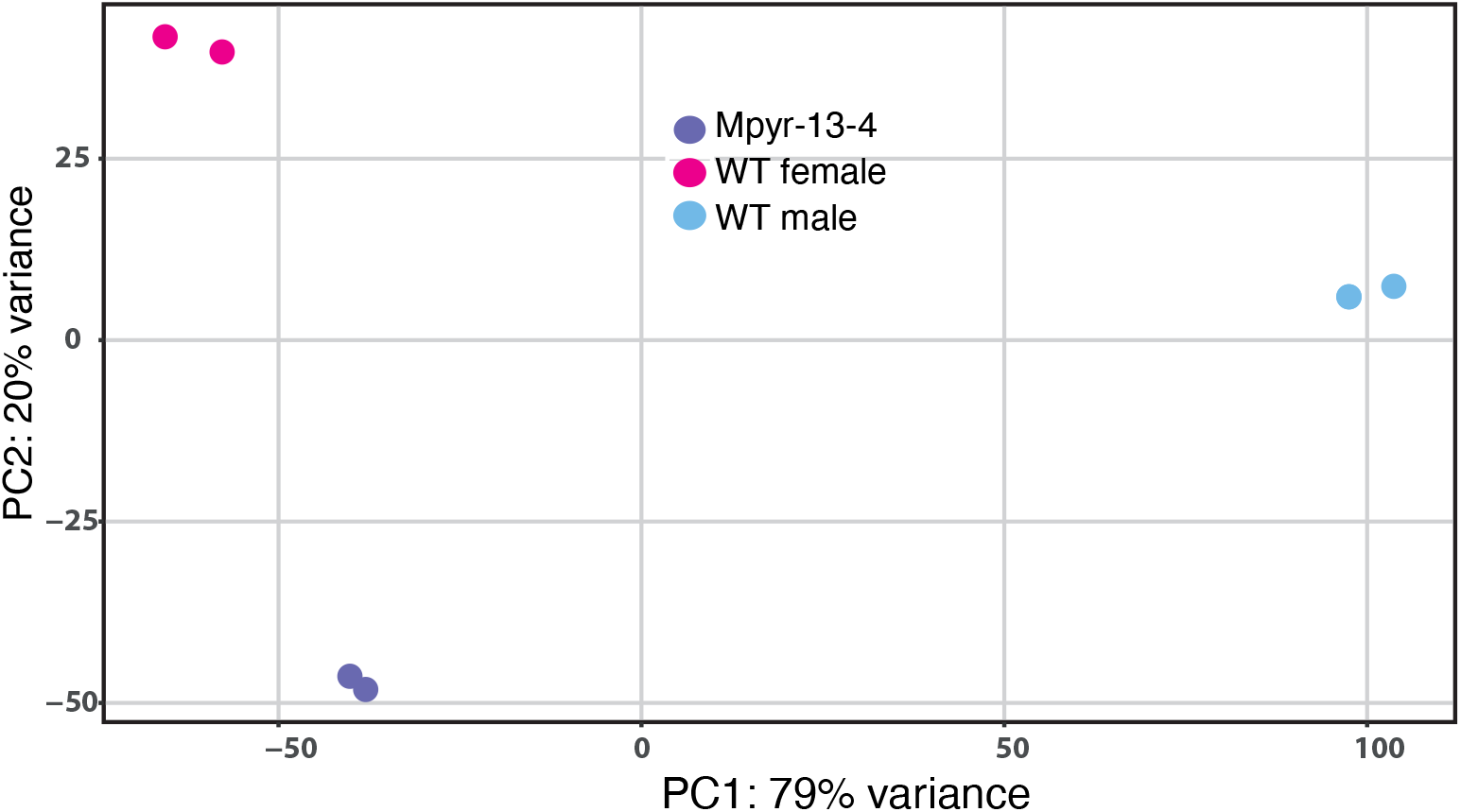
PCA was used to compare transcript abundance patterns across samples. The two dimensions represent 79% and 20% of the variance.

**Figure S4.**
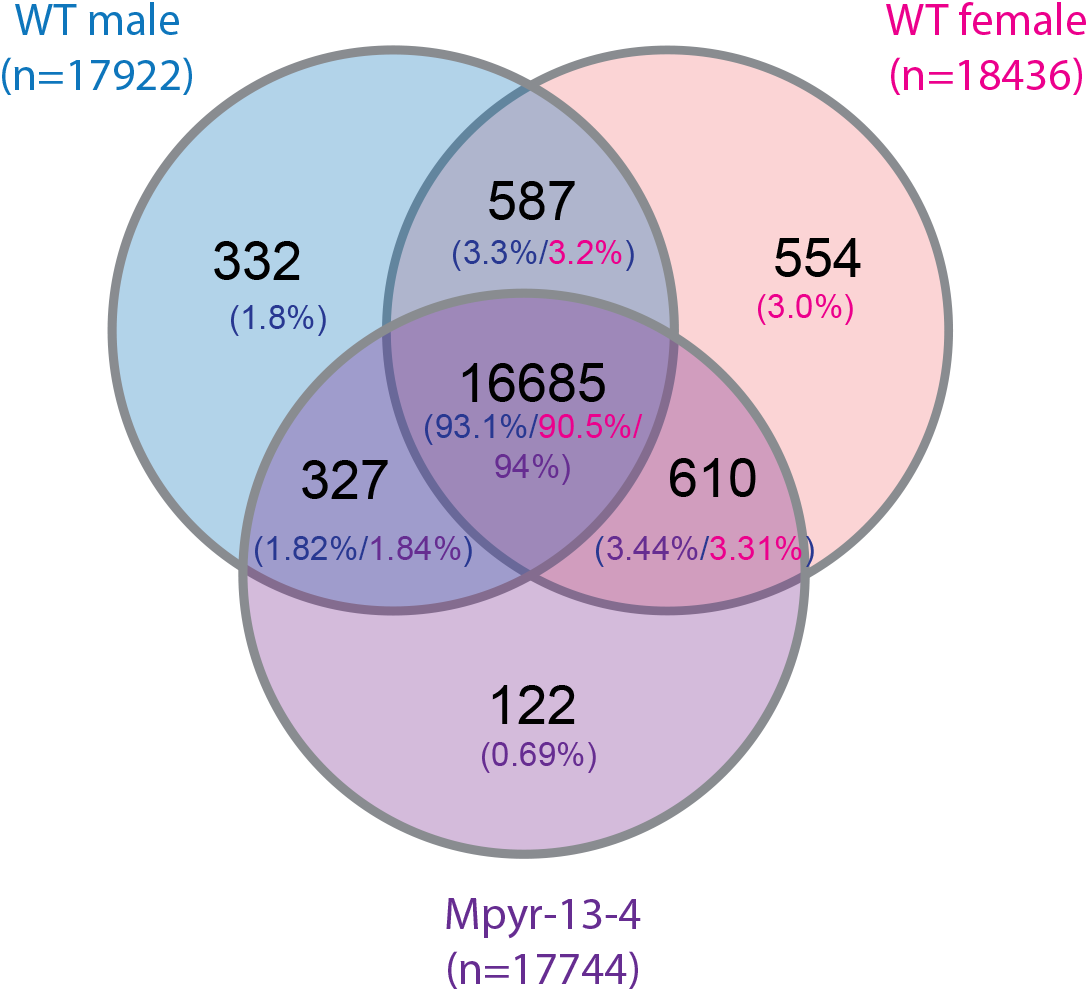
Venn diagram showing the sets of expressed genes (TPM>5th percentile) in wild-type male, wild-type female and variant Mpyr-13-4 lines and the overlap between the three sets.

**Figure S5.**
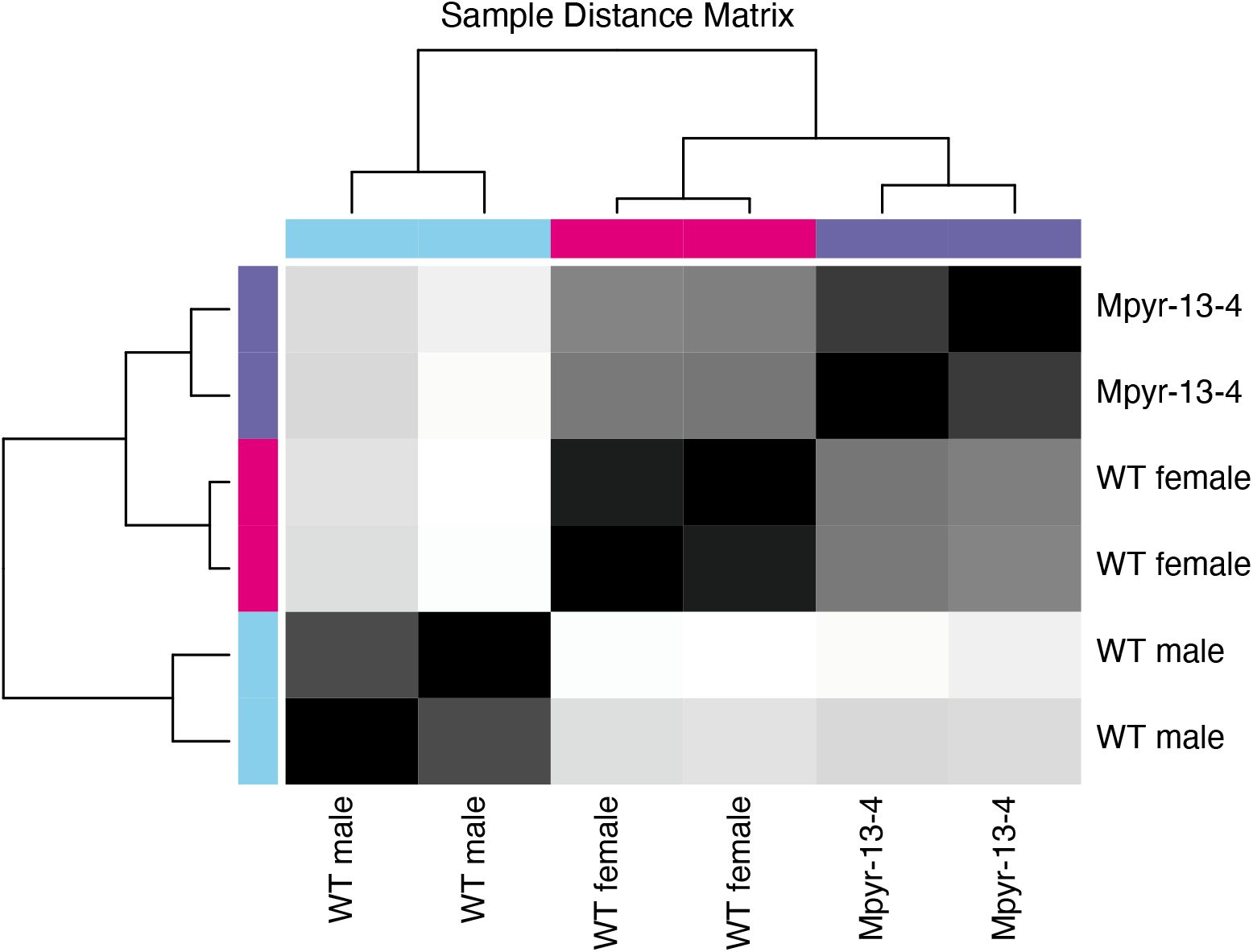
Expression heatmap of sample-to-sample distances on the matrix of variance-stabilized data for overall gene expression. Darker colors indicate more similar expression (color key is in arbitrary units). Clustering (top) demonstrates that the variant and female samples are very similar to each other but show complete separation from the male samples.

**Figure S6.**
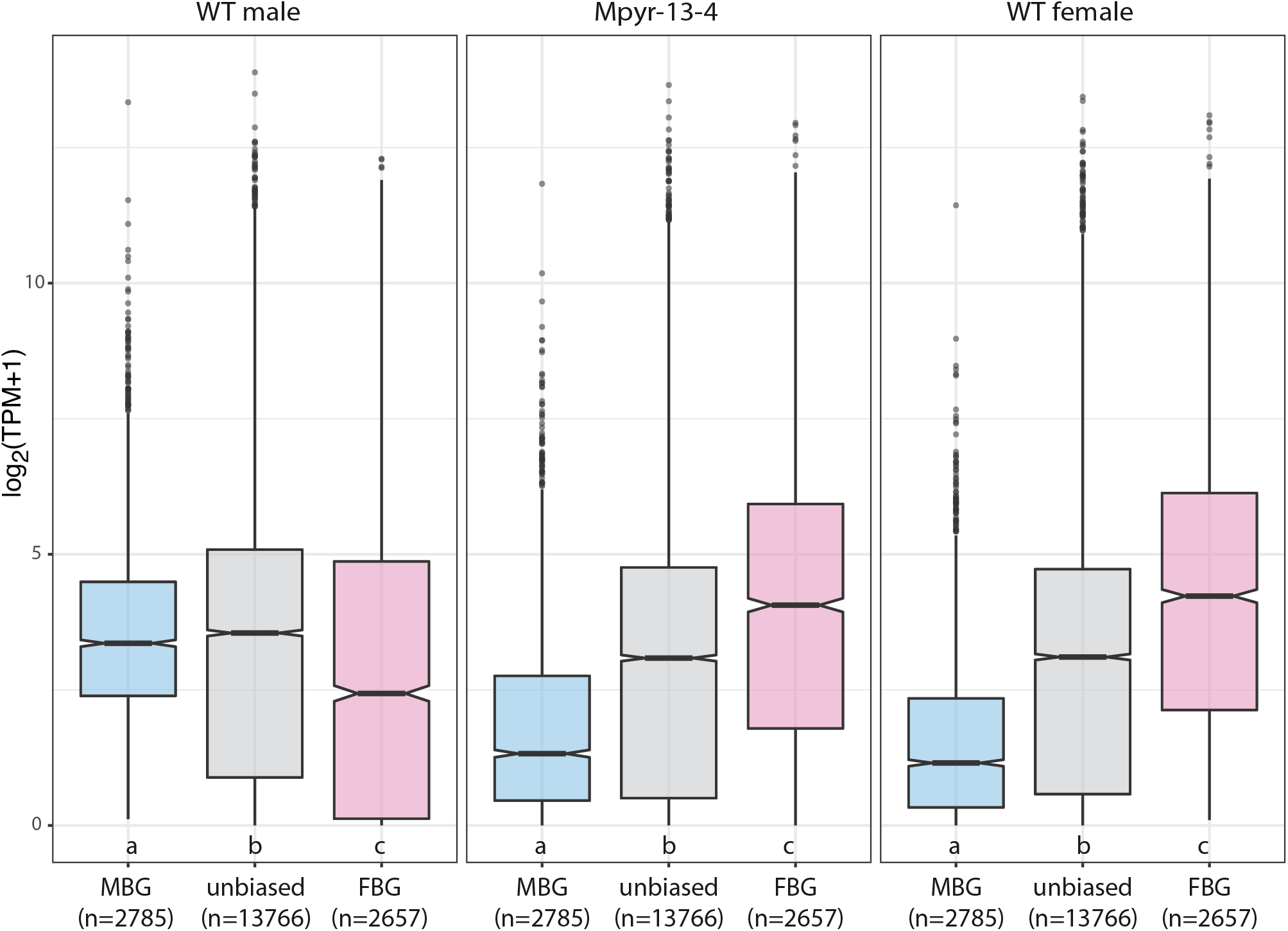
Levels of expression (log2TPM+1) of sex-biased genes in wild-type males, Mpyr-13-4 and wild-type females. MBG: male-biased genes; FBG: female-biased genes. Significant differences (Wilcoxon test, p<0.01) are indicated with different letters below the plots.

**Figure S7.**
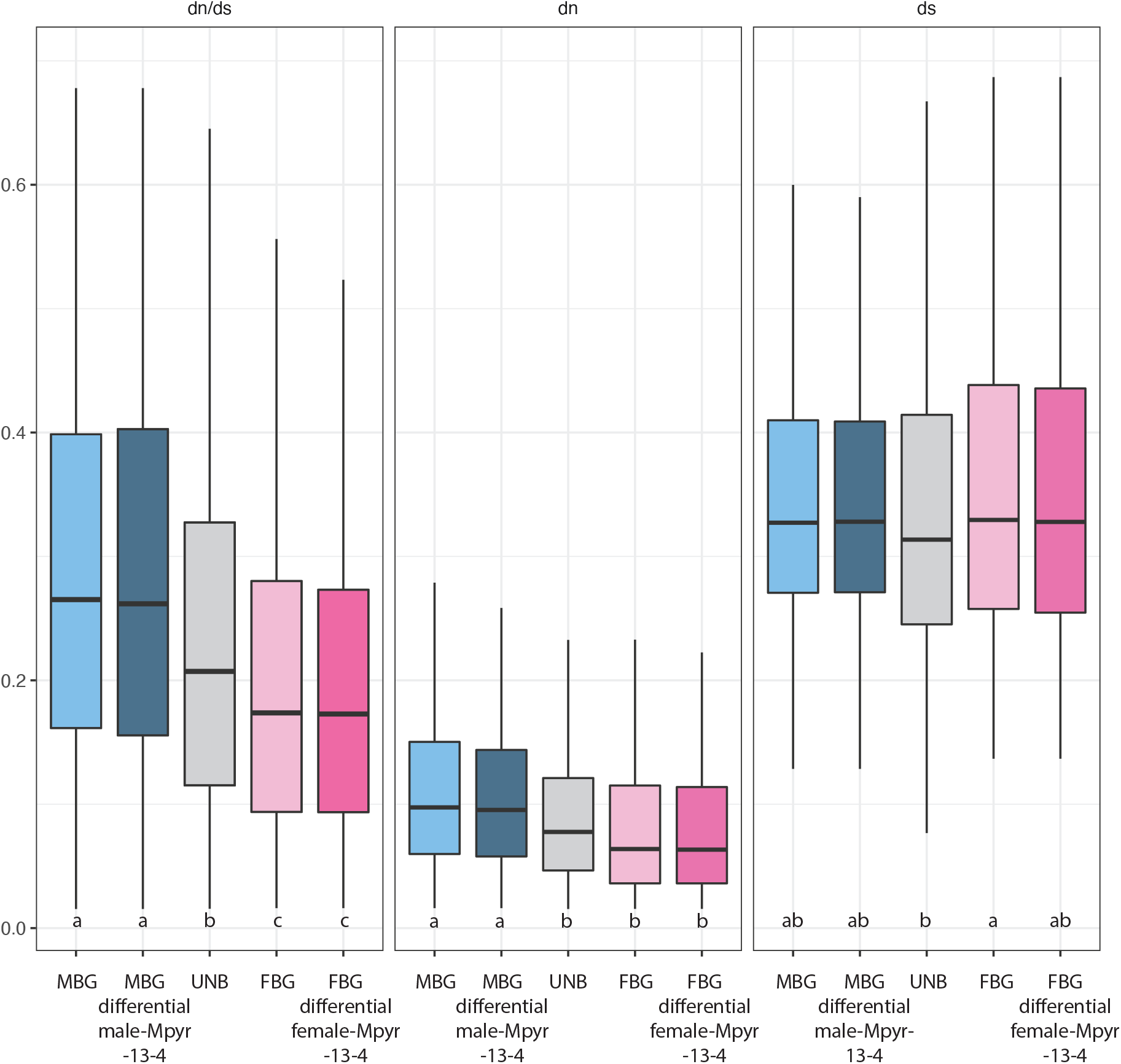
Rates of evolution of sex-biased genes that were differentially expressed in Mpyr-13-4 compared with wild-type males or wild-type females.

**Figure S8.**
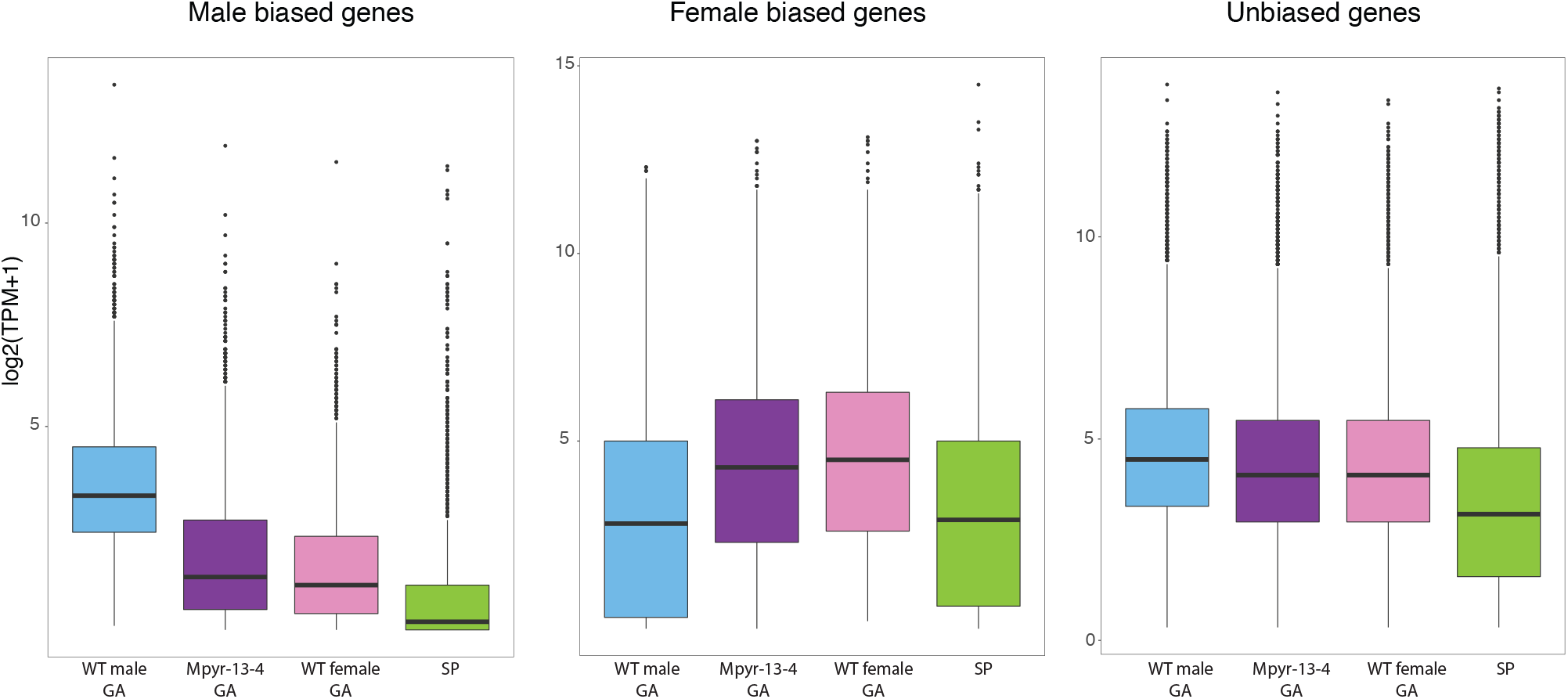
Expression of male-biased and female-biased genes (log_2_TPM) in gametophyte versus diploid sporophyte (SP) stage of development.

## Supplemental Tables

**Table S1**. List of strains used and genome and transcriptome assembly statistics.

**Table S2**. Number of biased genes from DEseq2 analysis and categories of sex-biased genes with different levels of FC between the three samples. Only genes with TPM>5th percentile in at least one of the samples were considered for the analysis.

**Table S3**. Differential expression levels (DEseq2, FC>2, padj<0.05) between wild-type male, wild-type female and Mpyr-13-4 samples.

**Table S4**. Gene expression (measured as TPM) in wild-type males, wild-type females and Mpyr-13-4 variant strain.

**Table S5**. Gene ontology terms significantly enriched among biased genes (Fisher exact test, FDR<5%).

## Acknowledgements

We thank Dominique Marie for help with the flow cytometry analysis, the Institut Français de Bioinformatique and the Roscoff Analysis and Bioinformatics for Marine Science platform ABiMS (http://abims.sb-roscoff.fr) for providing computing and data storage resources. This work was supported by the CNRS, Sorbonne Université and ERC grants to S.M.C. (grant agreement 638240 and 864038).

